# Analysis in the frequency domain of multicomponent oscillatory modes of the human electroencephalogram extracted with multivariate empirical mode decomposition

**DOI:** 10.1101/2020.06.06.138065

**Authors:** Eduardo Arrufat-Pié, Mario Estévez-Báez, José Mario Estévez-Carreras, Calixto Machado Curbelo, Gerry Leisman, Carlos Beltrán León

## Abstract

Considering the properties of the empirical mode decomposition to extract from a signal its natural oscillatory components known as intrinsic mode functions (IMFs), the spectral analysis of these IMFs could provide a novel alternative for the quantitative EEG analysis without a priori establish more or less arbitrary band limits. This approach has begun to be used in the last years for studies of EEG records of patients included in database repositories or including a low number of individuals or of limited EEG leads, but a detailed study in healthy humans has not yet been reported. Therefore, in this study the aims were to explore and describe the main spectral indices of the IMFs of the EEG in healthy humans using a method based on the FFT and another on the Hilbert-Huang transform (HHT). The EEG of 34 healthy volunteers was recorded and decomposed using a recently developed multivariate empirical mode decomposition algorithm. Extracted IMFs were submitted to spectral analysis with, and the results were compared with an ANOVA test. The first six decomposed IMFs from the EEG showed frequency values in the range of the classical bands of the EEG (1.5 to 56 Hz). Both methods showed in general similar results for mean weighted frequencies and estimations of power spectral density, although the HHT is recommended because of its better frequency resolution. It was shown the presence of the mode-mixing problem producing a slight overlapping of spectral frequencies mainly between the IMF3 and IMF4 modes.

## 1. Introduction

Brain bioelectric activity shows oscillations ranging from very slow frequencies with periods of several minutes or seconds to ultrahigh frequencies from 200 to 600 Hz (Hughes et al. 2011; Malekzadeh-Arasteh et al. 2020; van Putten et al. 2015; Vanhatalo et al. 2005). It has been found that several frequency ranges, generally known as bands, are associated with particular physiological functional states and in some cases with brain pathologies. For technological reasons associated with limitations and constraints of bio-amplifiers, the range from 0.5 to 30 Hz of the electroencephalogram (EEG) became the focus of the investigations since the seminal publication of Hans Berger (Berger 1929), reporting the presence of EEG oscillations recorded from the human scalp named by him as alpha and beta rhythms. Later, a definite physiological or pathological significance was associated to other bands (theta, delta, and gamma), and multiple rhythms and EEG features were described. With the development of improved bio-amplifiers, EEG oscillations with frequency higher to 30 Hz have been studied, conforming the so called gamma band, of high interest for cognitive and psychiatric studies (Amo et al. 2017; Park et al. 2014). A well established and routinely measured category of ultrafast EEG signals at 500-1000 Hz range are the brain stem evoked potentials. Intimately related to the study of the EEG frequency bands and rhythms, has evolved in parallel the study of the source of these oscillations in different cortical and subcortical brain structures (Buzsaki 2006; Buzsaki, Watson 2012; Smyk, van Luijtelaar 2020; Soltani Zangbar et al. 2020). Different studies have shown that when the frequency bands are arranged in increasing order of their center frequencies, can be observed some degree of overlapping of the oscillations of adjacent bands, and that the natural logarithm of these frequencies form a linearly increasing continuum (Buzsaki 2006; Penttonen, Buzsaki 2003).

A cardinal tool to the study of brain rhythms was the development of the FFT algorithm (Cooley, Tukey 1965) that was incorporated to the spectral analysis of the EEG and continues to be used nowadays with several improvements. However, the EEG dynamics shows nonlinearity and non-stationarity (Noshadi et al. 2014; Abdulhay et al. 2017; Alegre-Cortes et al. 2016; Soler et al. 2020) and strictly speaking, the FFT methods are limited to the study of linear systems. Therefore, in some conditions, the interpretation of the results obtained through the FFT could be meaningless.

A novel fully data-driven method (Huang et al. 1998) for the analysis of non-Gaussian, nonlinear and non-stationary signals, the empirical mode decomposition (EMD), followed by the Hilbert transform of the extracted modal components with the EMD, known also as the Hilbert-Huang transform method, has been introduced for the study of EEG signals (Al-Subari et al. 2015; Al-Subari et al. 2016; Carella et al. 2018; Chatterjee 2019; Chuang et al. 2019; Dinares-Ferran et al. 2018; Estevez-Baez et al. 2017a, b, c; Hansen et al. 2019; Hassan, Bhuiyan 2017; Hou et al. 2018; Javed et al. 2019; Rahman, Fattah 2017). The modal components extracted using the EMD can be considered as an adaptive spectral band (Noshadi et al. 2014). In the last years the study of the spectral frequency and power content of the oscillatory modes of the EEG extracted through EMD have been applied with varied aims, but in general these studies have used EEG recordings of patients from repositories of clinical databases, or have included a low number of individuals, or of EEG leads, and to the best of our knowledge the detailed study in healthy humans has not been reported. Besides, given the properties of the EMD to decompose a signal into its natural oscillatory components (Schiecke et al. 2019; Schiecke et al. 2015) it should be possible to explore the natural spectrum of the EEG without a priory established strict limits for the EEG bands. Therefore, in this study the aims have been to explore and describe the spectral frequency and power content of different component oscillatory modes of the EEG in healthy humans using two particular methods, to compare them, and to comment the possible implications for further studies.

## 2. Methods

### 2.1 Participants and general experimental profile

Thirty-four healthy right-handed volunteers were included in this study. Demographic and some vital indices of the participants are shown in Table 1.

**Table 1.** Demographic data and some vital signs of the thirty-four healthy volunteers.

| Gender | n | Age<br>(m ± sd) [range] | RR<br>(m ± sd) | HR<br>(m ± sd) | SBP<br>(m ± sd) | DBP<br>(m ± sd) |
| --- | --- | --- | --- | --- | --- | --- |
| Females | 18 | 33.59 ± 11.4 [18.7-57.6] | 15.67 ± 1.5 | 68.22 ± 7.7 | 114.17 ± 17.3 | 70.56 ± 11.1 |
| Males | 16 | 38.73 ± 15.6 [19.8-69.0] | 15.88 ± 2.0 | 68.88 ± 11.0 | 116.38 ± 10.3 | 74.94 ± 5.5 |
| Total | 34 | 36.01 ± 13.6 [18.7-69.0] | 15.76 ± 1.7 | 68.53 ± 9.3 | 115.21 ± 14.2 | 72.62 ± 9.1 |

Age, decimal age; n, number of subjects, m, mean value, sd, standard deviation RR, respiratory rate; HR, heart rate; SBP, systolic blood pressure; DBP, diastolic blood pressure

Inclusion criteria included an age over 18 year’s old, and voluntary participation. Were excluded individuals who reported any disease or health condition which could modify the EEG, or with a previous event of affected awareness without a clear diagnosis, and those who reported any toxic habit like alcoholism, smokers or psychotropic drugs consumption. The Ethics Committee of the Institute of Neurology and Neurosurgery approved this research and the participants gave their informed written consent to participate in the study. Volunteers were studied in a laboratory with controlled temperature from 22 to 24 °C, with noise attenuation and dimmed lights. A trained technician was present during the recording sessions, as well as one of us (EAP).

### 2.2 EEG recordings

EEG was recorded from 19 standard locations over the scalp according to the 10–20 system: Fp1, Fp2, F3, F4, C3, C4, P3, P4, O1, O2, F7, F8, T3, T4, T5, T6, Fz, Cz, and Pz. Silver scalp electrodes were fixed, after a careful cleaning of the skin, using a conductor paste, and connected to the input box of the digital EEG system (Medicid-05, Neuronic, S.A.). Monopolar leads were employed, using linked ears as a reference. EEG technical parameters were: gain 20,000, pass-band filters 0.5–70 Hz, “notch” filter at 60 Hz, noise level of 2 µV (root mean square), sampling frequency 200 Hz, A/D converter of 15 bits, and electrode–skin impedance never higher than 5 KΩ. Two of us (EAP and MEB) visually inspected the records to select continuous free of artifacts EEG segments with a total duration of no less than 60 seconds, which were later exported to an ASCII file, for further quantitative analysis.

### 2.3 EEG Pre-processing

A digital bandpass “FIR” filter created with Matlab using the function "desingnfilt.m" and the constrained least squares design method with cutoff frequencies fixed at 1 and 70 Hz was applied to the EEG 60-seconds segments digitally stored, using the “filtfilt.m” function to achieve a “zero-phase” distortion effect. Finally, to avoid outliers, the function “filloutliers.m” was applied using for filling the outliers a piecewise cubic spline interpolation method.

### 2.4 Empirical mode decomposition (EMD)

EMD is a fully data-driven method for decomposing a time series into AM/FM components which reflects its natural oscillations. In EMD the original signal x(t) is decomposed as a linear combination of data-driven set of basic functions known as intrinsic mode functions (IMFs) denoted by m_k_ (t), k = 1…K, where K is the total number of IMFs generated for the signal. EMD algorithm extracts the IMFs using an iterative technique called “sifting”.

IMFs have two basic properties:

1. In the whole data set, the number of extrema is equal to the number of zero-crossings or differ at most by one.
2. At any point, the mean value of the envelope defined by the local maxima and the envelope defined by the local minima is zero.

Once an IMF is obtained, it is subtracted from the original signal and the procedure is repeated on the remaining signal to obtain the next IMF. The process is stopped when the signal at the end of an iteration becomes a constant, a monotonic function, or a function containing only a single extremum.

The signal x(t) can then be represented as in the expression

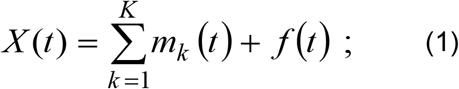

where m_k_ are the IMFs and f(t) is the final residual.

### 2.5 Multivariate empirical mode decomposition (MEMD)

Multivariate data, as it is the case of EEG signals, contain joint rotational modes whose coherent treatment is required for time-frequency estimation. In EMD, the local mean is calculated by taking an average of upper and lower envelopes, which in turn are obtained by interpolating between the local maxima and minima. However, generally, for multivariate signals, the local maxima and minima may not be defined directly. Besides, the notion of ‘oscillatory modes’ defining an IMF is rather confusing for multivariate signals. MEMD propose to generate multiple n-dimensional envelopes by taking signal projections along different directions in n-dimensional spaces; these are then averaged to obtain the local mean.

This calculation of the local mean may be considered an approximation of the integral of all the envelopes along multiple directions in an n-dimensional space. As the direction vectors in n-dimensional spaces can be equivalently represented as points on the corresponding unit (n − 1) spheres, the problem of finding a suitable set of direction vectors can be treated as that of finding a uniform sampling scheme on an n-sphere. A convenient choice for a set of direction vectors can be to employ uniform angular sampling of a unit sphere in an n-dimensional hyper-spherical coordinate system.

A coordinate system in an n-dimensional Euclidean space can be defined to serve as a pointset (and the corresponding set of direction vectors) on an (n − 1) sphere. To generate a uniform pointset on an n-sphere it is used the concept of discrepancy. A convenient method for generating multidimensional ‘low-discrepancy’ sequences involves the family of Halton and Hammersley sequences.

Whereas complete MEMD algorithm is described elsewhere (Rehman, Mandic 2010; Ur Rehman et al. 2010) only a brief description for those unfamiliar with the technique is added in Table 2.

**Table 2.**
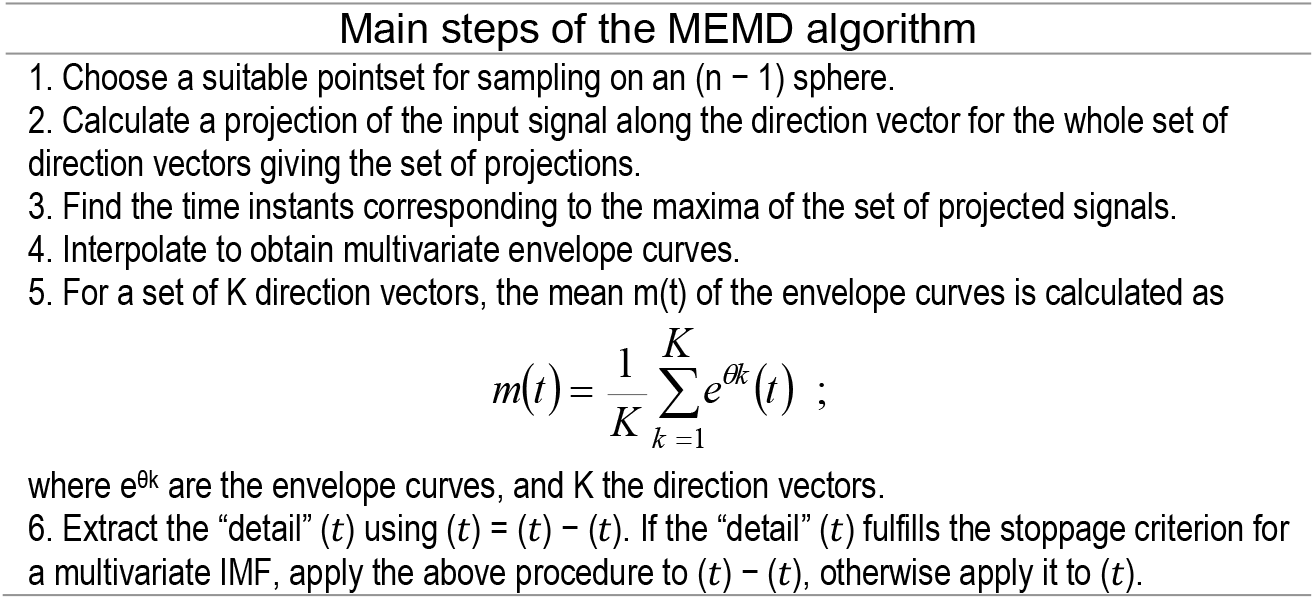
Algorithm of the multivariate empirical mode decomposition.

| Main steps of the MEMD algorithm |
| --- |
| 1. Choose a suitable pointset for sampling on an $(n - 1)$ sphere.<br>2. Calculate a projection of the input signal along the direction vector for the whole set of direction vectors giving the set of projections.<br>3. Find the time instants corresponding to the maxima of the set of projected signals.<br>4. Interpolate to obtain multivariate envelope curves.<br>5. For a set of $K$ direction vectors, the mean $m(t)$ of the envelope curves is calculated as $m(t) = \frac{1}{K} \sum_{k=1}^K e^{\theta_k}(t) ;$<br>where $e^{\theta_k}$ are the envelope curves, and $K$ the direction vectors.<br>6. Extract the "detail" $(t)$ using $(t) = (t) - (t)$. If the "detail" $(t)$ fulfills the stoppage criterion for a multivariate IMF, apply the above procedure to $(t) - (t)$, otherwise apply it to $(t)$. |

### 2.6 Adaptive-projection intrinsically transformed MEMD (APIT-MEMD)

Recently, it has been developed an extension to the MEMD algorithm to counteract for power imbalances and inter-channel correlations in multichannel data of the EEG (Hemakom et al. 2016). The main steps of this novel algorithm are included in Table 3.

**Table 3.** Algorithm of the adaptive-projection intrinsically transformed MEMD.

| Main steps of the APIT-MEMD algorithm |
| --- |
| 1. Given an $n$-variate input signal to each sifting operation $s(t)$, perform the eigendecomposition of the covariance, $C=E\{ss^T\} = V\Lambda V^T$, where $V=[v_1, v_2, \dots, v_n]$ is the eigenvector matrix, and $\Lambda=\text{diag}\{\lambda_1, \lambda_2, \dots, \lambda_n\}$ is the eigenvalue matrix, with the largest eigenvalue $\lambda_1$ corresponding to eigenvector $v_1$ (first principal component).<br>2. Construct a vector pointing in the diametrically opposite direction to the first principal component vector in $n$-dimensional space.<br>3. Uniformly sample an $(n - 1)$ sphere using the Hammersley sequence to generate a set of $K$ direction vectors<br>4. Calculate the Euclidean distances from each of the uniform direction vectors to $v_1$.<br>5. Half of all the uniform projection vectors $X_{v1}^{\theta_k}$ nearest to $v_1$ are relocated using the expression $\hat{X}_{v1}^{\theta_k} = \frac{X_{v1}^{\theta_k} + \alpha_{v1}}{\vert X_{v1}^{\theta_k} + \alpha_{v1}\vert };$<br>where $\alpha$ parameter is used to control the density of the relocated vectors.<br>6. The other half of all the uniform projection vectors, $X_{v01}^{\theta_k}$ nearest to $v_{01}$ are relocated using $\hat{X}_{v1}^{\theta_k} = \frac{X_{v01}^{\theta_k} + \alpha_{v01}}{\vert X_{v01}^{\theta_k} + \alpha_{v01}\vert };$<br>where $\alpha$ is used to control the density of the relocated vectors.<br>7. Perform local mean estimation according to the conventional MEMD algorithm, using the adaptive direction vectors $\hat{X}_{v1}^{\theta_k}$ and $X_{v01}^{\theta_k}$ |

In this study we used the APIT-MEMD algorithm for the EEG decomposition, with an alpha parameter of 0.35 and the number of directions was of 128. For calculations in the frequency domain of EEG signals were selected the first 6 extracted IMFs, that contain the spectral range from 1 to 70 Hz as shown by other authors, and our own previous experience (Amo et al. 2017; Carella et al. 2018; Chen et al. 2016; Chen et al. 2017; Schiecke et al. 2015; Schiecke et al. 2019; Tsai et al. 2016; Zheng, Xu 2019; Zhuang et al. 2017; Estevez-Baez et al. 2017a, c). The implementation of the MEMD and the APIT-MEMD algorithms used in this study was based on software resources publicly available at the web site http://www.commsp.ee.ic.ac.uk/~mandic/research/emd.htm.

### 2.7 Estimation of the PSD of the IMFs using the FFT method, and spectral indices (FFT)

The power spectrum was calculated for each extracted IMF of every EEG lead using the Welch periodogram method, with the Matlab function “pwelch.m”, with a window of 512 samples, with a 50% of overlapping, a Hamming window, and 2000 samples for the FFT. The spectral resolution was 0.1 Hz.

The absolute spectral power of each IMF was calculated for the whole time duration of the EEG segments from which were extracted the corresponding IMFs (60 s) following the expression:

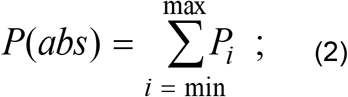

where P_i_ are the discrete power values of the spectra of the IMFs, ‘min’ represents the lower limit value of the spectral range to consider (1 Hz) and ‘max’ the upper limit (70 Hz in this study).

The spectral relative power of each IMF expressed in normalized units (%) was calculated as:

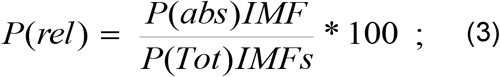

where P(abs)IMF is the absolute power of the IMF, and P(Tot)IMFs is the sum of the absolute power of the six Welch IMF spectra.

The mean weighted frequency (Xie, Wang 2006) of each IMF spectra was calculated using the expression:

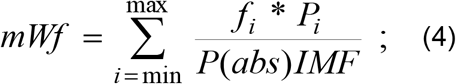

where f_i_ discrete spectral frequencies, P_i_ corresponding discrete spectral powers, and P(abs)IMF is the absolute power calculated for the spectrum of the corresponding IMF, ‘min’ is the lower value of the spectral range (I Hz), and ‘max’ the upper limit (70 Hz).

### 2.8 Estimation of the PSD of the IMFs using the HHT method and spectral indices (HHT)

The Hilbert transform was applied to each extracted IMF of every EEG lead to create analytic functions, referred here as Z(t), that can be written as

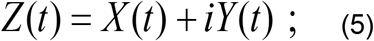

where X(t) is the input time series (the IMF in this case), and iY(t) is the Hilbert transform of X(t). The instantaneous values of power and frequency can then be obtained using the following expressions:

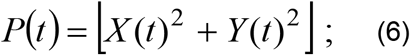

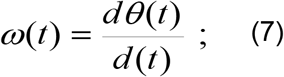

where P(t) is the instantaneous energy power and ω(t) is the instantaneous frequency (Estevez-Baez et al. 2019; Estevez-Baez et al. 2018).

Spectral indices were then calculated for the 60-seconds EEG segments, from which were extracted the corresponding IMFs. The absolute spectral power of each IMF was calculated using the expression:

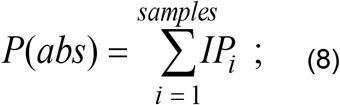

where IP_i_ are the consecutive values of instantaneous power obtained from each sample of the input signal, that in in this case was the IMF with 12000 samples (60 seconds of EEG recording at 200 Hz of sampling frequency).

The spectral relative power of the IMFs expressed in normalized units (%) was calculated as:

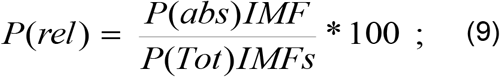

where P(abs)IMF is the absolute power of the IMF, and P(Tot)IMFs is the sum of the absolute power of the six IMFs.

The mean weighted frequency of each IMF, following also the recommendation of (Xie, Wang 2006), was calculated using the expression:

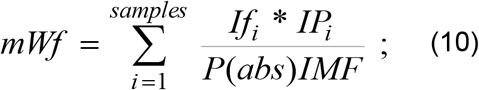

where If_i_ consecutive instantaneous frequencies, IP_i_ corresponding values of instantaneous powers, and P(abs)IMF is the absolute power calculated for the corresponding IMF, and samples in this case were 12000.

### 2.9 Statistical analysis

All results were calculated in this study using custom tailored scripts and functions developed by two of us (MEB and JMEC) using Matlab R2019b (version 9.7.0.1190202, The MathWorks, Inc.). The grand average method was used to calculate the distribution of the instantaneous frequencies calculated for the IMFs in several EEG leads of interest, and of the power spectra of the IMFs calculated with the FFT method. It was explored for all the calculated indices the normality of their distribution using the Shapiro-Wilks, Lilliefors, and the Kolmogorov-Smirnov tests, and transformed when necessary to achieve a normal distribution before the statistical comparisons. An ANOVA test of repeated measures was carried out for the comparison of the values obtained using both methods of spectral analysis using the statistical commercial package Statistica 10 (StatSoft, Inc. (2011).

## 3. Results

The multivariate empirical mode decomposition using the APIT-MEMD algorithm showed the same number of decomposed IMFs for all the EEG leads in each subject. The decomposition process is shown graphically for two symmetrical (C3 and C4) EEG leads in one of the volunteers (See Figure 1).

**Figure 1.**
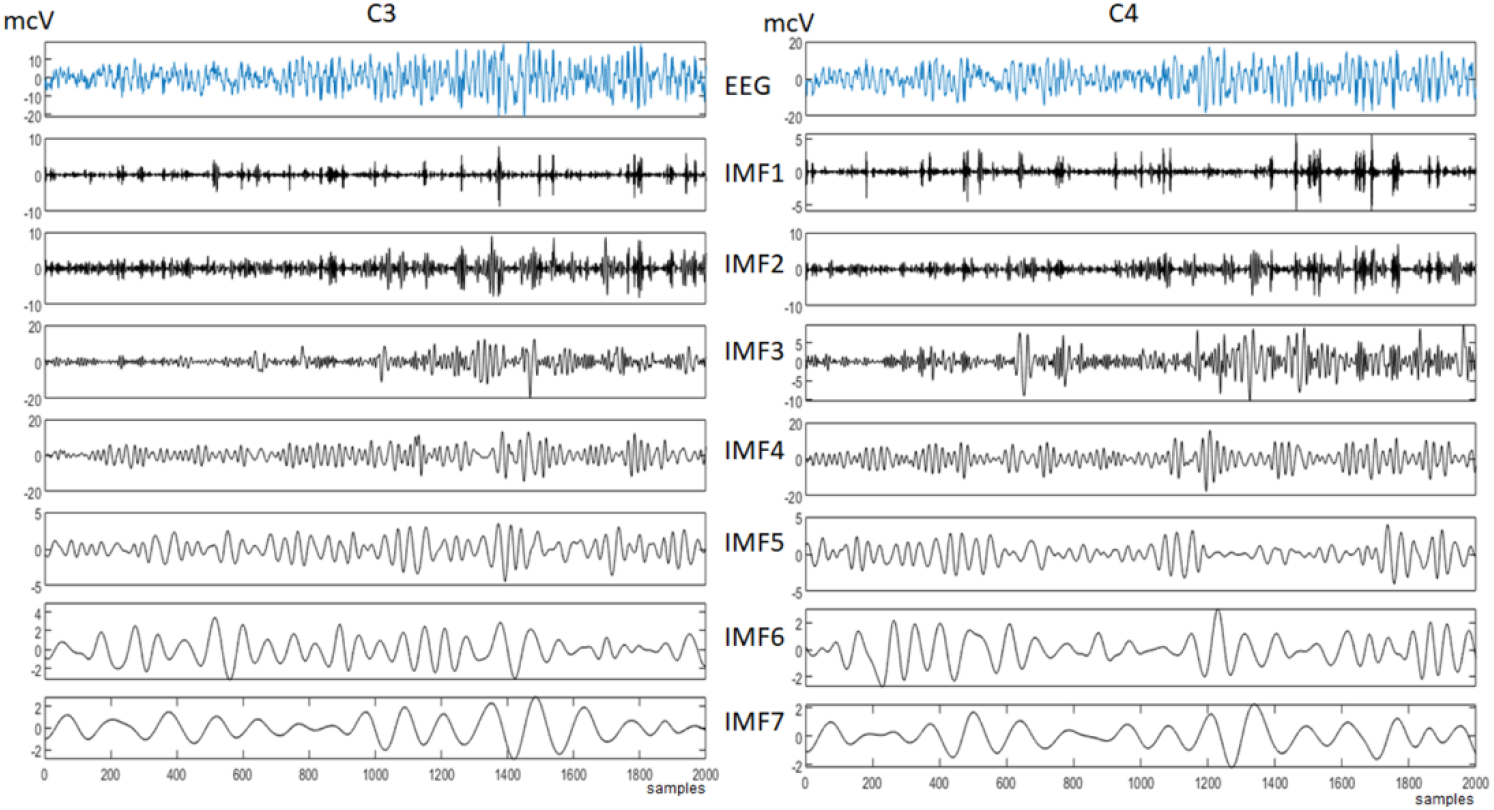
Decomposition of EEG leads C3 and C4 using the APIT-MEMD algorithm in one representative healthy volunteer. The original segment of the EEG is shown in the top diagram of each panel and the corresponding IMFs are shown in descending order of frequency content. The IMF7 has been included in the figure although it was not considered in this study because its mean frequency for the whole group (0.45 Hz) was under the lower limit of the spectral range investigated (> 1 Hz).

The power spectra of the six IMFs showed that their magnitude was higher for some of them, particularly for the IMF3 and IMF4, and this issue hinders the visual inspection of the spectra of all the IMFs in a same diagram as can be observed in Figure 2 A and B. Alternatively, an individual IMF spectra representation can be used (See Figure 2 A’, and B’). However, no matter what kind of graphical method was used, it resulted evident for most of the spectra of the IMFs of all the EEG leads, the overlapping of spectral content between adjacent IMFs.

**Figure 2.**
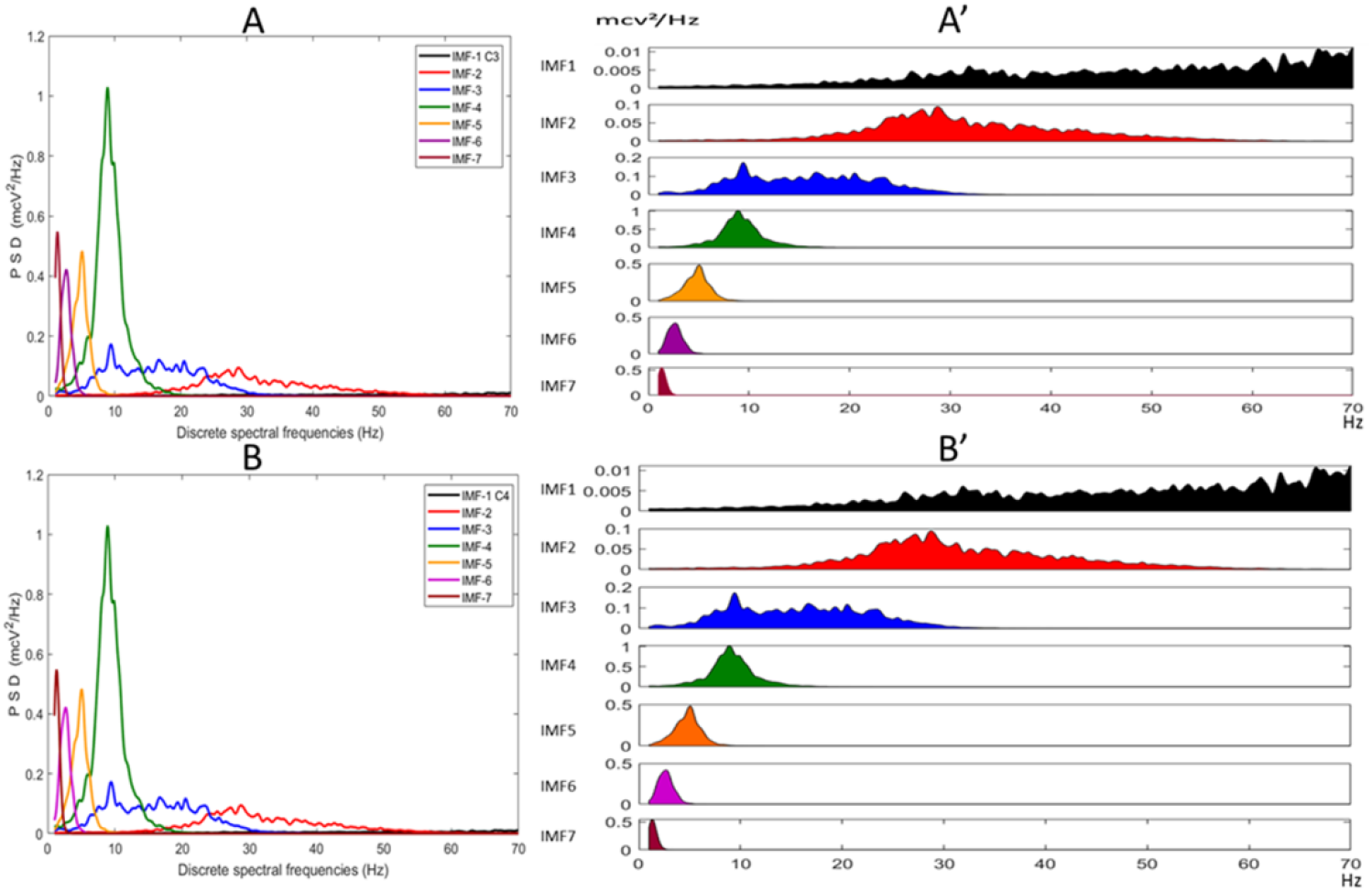
Power spectra of the IMFs extracted using the APIT-MEMD algorithm in a representative healthy volunteer. Panels A and B show the superimposed spectra corresponding to the EEG leads C3 and C4, while in panels A’ and B’ are shown the same power spectra of the individual using other format that avoids the effects of the different values of the amplitude of the spectra IMFs for the visual inspection. The IMF7 has been included in the figure although it was not considered in this study because its mean frequency for the whole group (0.45 Hz) was under the lower limit of the spectral range investigated (> 1 Hz).

The sequence of consecutive instantaneous values of frequency and power obtained after the Hilbert transformation of the IMFs in one of the volunteers, can be observed for a 2000 samples segment (10 s) in Figure 3.

**Figure 3.**
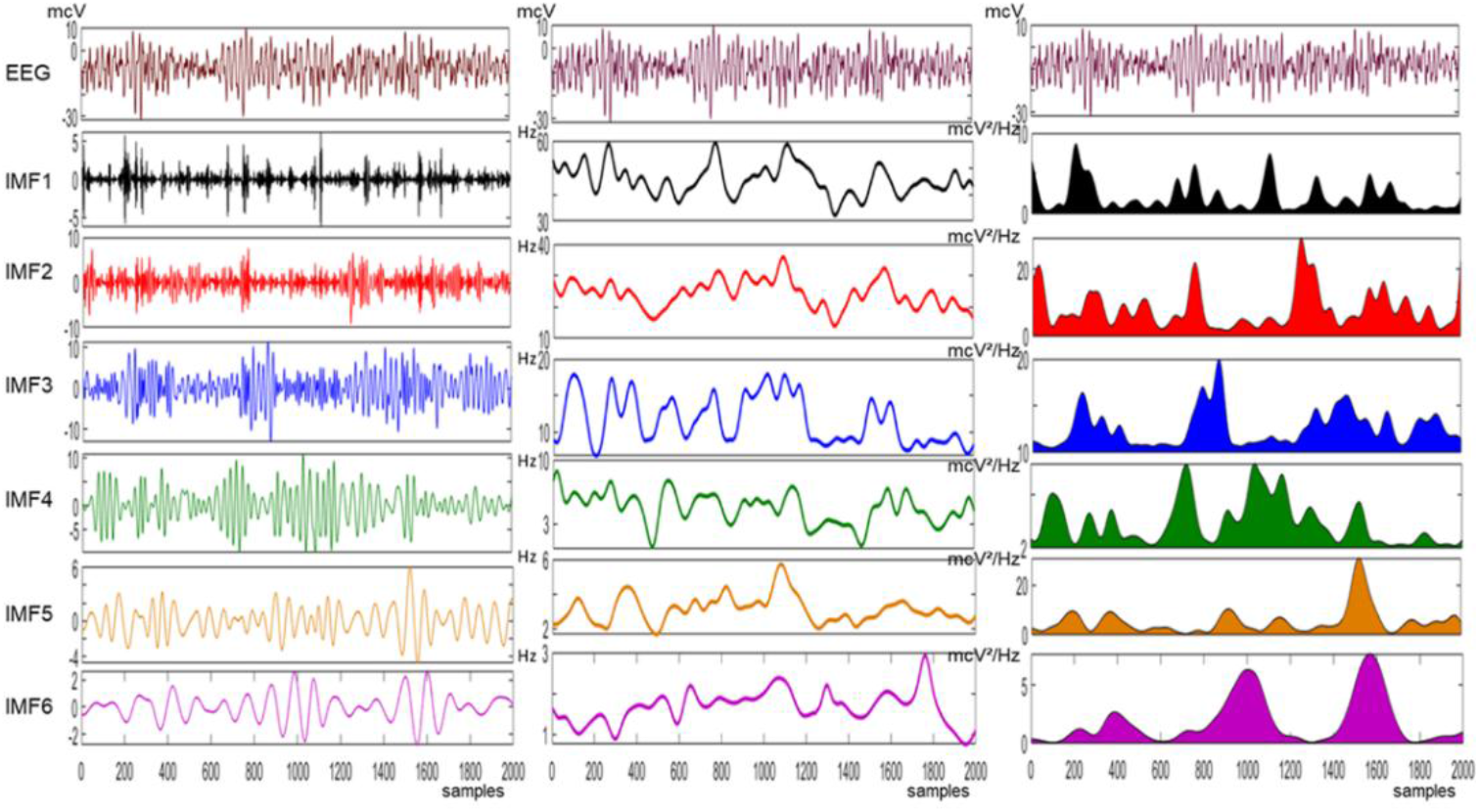
Collage showing in the left panel the first 6 IMFs decomposed from the EEG segment of a healthy volunteer using APIT-MEMD. The central panel depicts the instantaneous values of spectral frequency of the corresponding IMFs, and in the right panel are shown the instantaneous values of spectral power obtained using the HHT transform.

The method of grand averages showed that the overlapping of spectral content between IMFs, observed on Figure 2A and B for an individual, was also found for the whole group of healthy volunteers (See Figure 4 B, C). The grand average of the distributions of the instantaneous frequencies of the IMFs showed in general a normal distribution for the EEG leads. An illustration of this fact is shown related to the results obtained for the EEG lead C3 (Figure 4A). This approach also showed the presence of similar frequencies in different IMFs, but the degree of the overlapping was not as marked as when analyzed using the FFT spectra of the IMFs for the same EEG lead (Figure 4 B and C).

**Figure 4.**
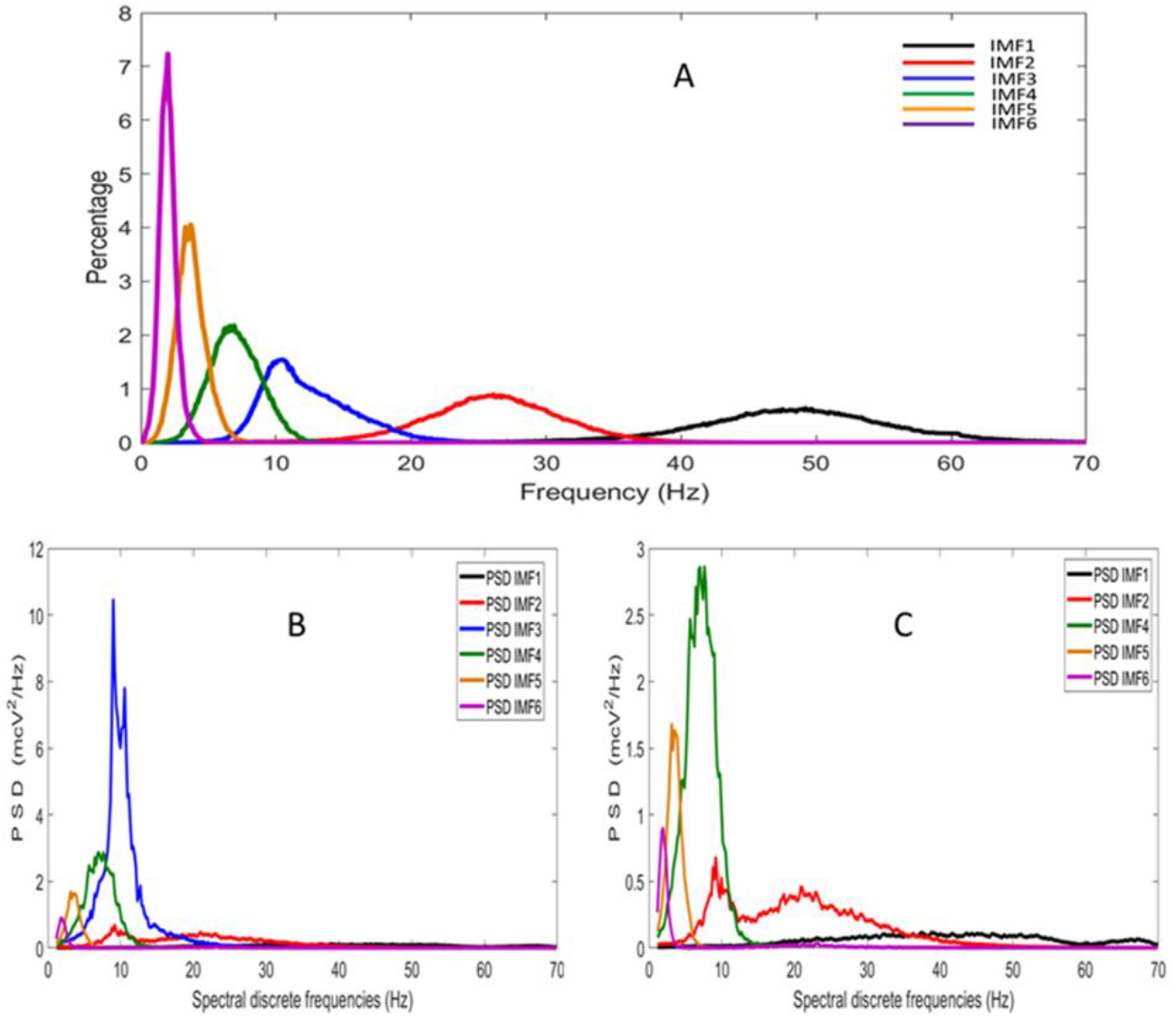
Panel labeled A shows the grand average of distributions of the instantaneous frequency for the first 6 IMFs decomposed from the EEG of 34 healthy volunteers using the APIT-MEMD algorithm and the Hilbert-Huang transform. Panel B depicts the grand averages of the FFT spectra of the first six IMFs, while on Panel C, has been excluded the average spectrum corresponding to the IMF3 to facilitate the visual inspection of IMFs with lower PSD values (IMF1 and IMF2). Results correspond to the calculations for EEG-lead C3.

The mean weighted spectral frequencies calculated with the two methods resulted in values that significantly differed. The HHT method showed significantly higher values for all the EEG leads of the IMFs 1 to IMF3, and lower values for the IMF6. There were not found differences for the values calculated for the IMF4 and IMF5 (See Table 4). However, the values corresponding to the estimations of the PSD in normalized units (%) only showed isolated differences in some EEG leads (See Table 5).

**Table 4.** Mean weighted frequency values (Hz) calculated from the intrinsic mode functions extracted from the EEG of thirty-four healthy volunteers using two spectral methods.

| Leads | Method | IMF1 | IMF2 | IMF3 | IMF4 | IMF5 | IMF6 |
| --- | --- | --- | --- | --- | --- | --- | --- |
| Fp1 | FFT | 44.36 ± 3.8 | 26.08 ± 1.8 | 12.53 ± 1.5 | 8.47 ± 0.9 | 4.25 ± 0.4 | 2.29 ± 0.3 |
|  | HHT | 49.85 ± 2.9 ↑ | 29.23 ± 1.9 ↑ | 13.64 ± 1.7 ↑ | 8.46 ± 0.9 | 4.28 ± 0.4 | 2.18 ± 0.3 ↓ |
| Fp2 | FFT | 44.23 ± 4.0 | 26.12 ± 1.8 | 12.51 ± 1.5 | 8.44 ± 0.9 | 4.24 ± 0.4 | 2.26 ± 0.2 |
|  | HHT | 49.81 ± 3.0 ↑ | 29.28 ± 1.8 ↑ | 13.59 ± 1.7 ↑ | 8.39 ± 1.0 | 4.24 ± 0.4 | 2.15 ± 0.3 ↓ |
| F3 | FFT | 43.95 ± 3.7 | 25.99 ± 1.9 | 12.53 ± 1.5 | 8.49 ± 0.9 | 4.29 ± 0.3 | 2.27 ± 0.2 |
|  | HHT | 49.64 ± 2.6 ↑ | 29.24 ± 2.0 ↑ | 13.65 ± 1.7 ↑ | 8.41 ± 1.0 | 4.30 ± 0.3 | 2.19 ± 0.3 ↓ |
| F4 | FFT | 43.77 ± 3.4 | 25.95 ± 1.8 | 12.50 ± 1.5 | 8.43 ± 0.9 | 4.24 ± 0.4 | 2.27 ± 0.2 |
|  | HHT | 49.51 ± 2.4 ↑ | 29.30 ± 1.9 ↑ | 13.67 ± 1.7 ↑ | 8.34 ± 0.9 | 4.23 ± 0.4 | 2.19 ± 0.3 ↓ |
| C3 | FFT | 43.52 ± 3.7 | 25.80 ± 1.8 | 12.33 ± 1.4 | 8.68 ± 0.9 | 4.30 ± 0.4 | 2.26 ± 0.3 |
|  | HHT | 49.50 ± 2.6 ↑ | 29.40 ± 1.9 ↑ | 13.45 ± 1.6 ↑ | 8.71 ± 1.0 | 4.33 ± 0.4 | 2.18 ± 0.3 ↓ |
| C4 | FFT | 43.71 ± 4.0 | 25.81 ± 2.0 | 12.24 ± 1.3 | 8.70 ± 0.9 | 4.29 ± 0.4 | 2.27 ± 0.2 |
|  | HHT | 49.83 ± 2.8 ↑ | 29.37 ± 2.1 ↑ | 13.44 ± 1.5 ↑ | 8.72 ± 1.0 | 4.28 ± 0.4 | 2.18 ± 0.3 ↓ |
| P3 | FFT | 43.45 ± 4.0 | 25.78 ± 1.8 | 12.36 ± 1.3 | 8.99 ± 0.9 | 4.35 ± 0.4 | 2.27 ± 0.2 |
|  | HHT | 49.57 ± 2.6 ↑ | 29.38 ± 1.9 ↑ | 13.53 ± 1.6 ↑ | 8.98 ± 1.1 | 4.35 ± 0.4 | 2.17 ± 0.3 ↓ |
| P4 | FFT | 43.49 ± 3.8 | 25.66 ± 1.8 | 12.30 ± 1.2 | 8.96 ± 1.0 | 4.31 ± 0.4 | 2.30 ± 0.3 |
|  | HHT | 49.57 ± 2.5 ↑ | 29.34 ± 2.0 ↑ | 13.53 ± 1.5 ↑ | 8.98 ± 1.2 | 4.29 ± 0.4 | 2.20 ± 0.3 ↓ |
| O1 | FFT | 43.62 ± 3.6 | 25.66 ± 1.7 | 12.10 ± 1.2 | 9.10 ± 1.1 | 4.34 ± 0.4 | 2.31 ± 0.2 |
|  | HHT | 49.54 ± 2.3 ↑ | 29.33 ± 1.7 ↑ | 13.28 ± 1.6 ↑ | 9.15 ± 1.2 | 4.35 ± 0.4 | 2.22 ± 0.3 ↓ |
| O2 | FFT | 43.43 ± 3.2 | 25.80 ± 1.6 | 12.00 ± 1.3 | 9.28 ± 1.1 | 4.37 ± 0.4 | 2.34 ± 0.3 |
|  | HHT | 49.30 ± 1.7 ↑ | 29.45 ± 1.7 ↑ | 13.16 ± 1.6 ↑ | 9.36 ± 1.3 | 4.32 ± 0.4 ↓ | 2.26 ± 0.3 ↓ |
| F7 | FFT | 43.86 ± 3.2 | 25.93 ± 1.8 | 12.55 ± 1.5 | 8.55 ± 0.9 | 4.27 ± 0.3 | 2.31 ± 0.2 |
|  | HHT | 49.37 ± 2.3 ↑ | 29.04 ± 1.8 ↑ | 13.68 ± 1.7 ↑ | 8.54 ± 1.0 | 4.23 ± 0.3 | 2.19 ± 0.3 ↓ |
| F8 | FFT | 44.07 ± 3.7 | 26.04 ± 1.9 | 12.46 ± 1.4 | 8.51 ± 0.9 | 4.23 ± 0.4 | 2.35 ± 0.2 |
|  | HHT | 49.64 ± 2.6 ↑ | 29.30 ± 1.9 ↑ | 13.56 ± 1.6 ↑ | 8.47 ± 1.0 | 4.23 ± 0.4 | 2.25 ± 0.3 ↓ |
| T3 | FFT | 43.84 ± 3.3 | 26.03 ± 1.9 | 12.56 ± 1.4 | 8.64 ± 0.9 | 4.31 ± 0.4 | 2.28 ± 0.2 |
|  | HHT | 49.40 ± 2.3 ↑ | 29.27 ± 1.8 ↑ | 13.66 ± 1.6 ↑ | 8.68 ± 1.1 | 4.28 ± 0.3 | 2.19 ± 0.3 ↓ |
| T4 | FFT | 43.91 ± 3.5 | 26.01 ± 2.0 | 12.44 ± 1.3 | 8.67 ± 1.0 | 4.33 ± 0.4 | 2.32 ± 0.3 |
|  | HHT | 49.49 ± 2.2 ↑ | 29.23 ± 1.8 ↑ | 13.54 ± 1.4 ↑ | 8.66 ± 1.1 | 4.31 ± 0.3 | 2.23 ± 0.3 ↓ |
| T5 | FFT | 43.99 ± 3.9 | 25.85 ± 1.8 | 12.40 ± 1.3 | 8.98 ± 1.0 | 4.35 ± 0.3 | 2.27 ± 0.3 |
|  | HHT | 49.67 ± 2.7 ↑ | 29.16 ± 1.6 ↑ | 13.56 ± 1.5 ↑ | 9.02 ± 1.2 | 4.30 ± 0.3 ↓ | 2.17 ± 0.3 ↓ |
| T6 | FFT | 43.71 ± 3.6 | 25.92 ± 1.7 | 12.41 ± 1.3 | 8.91 ± 1.1 | 4.40 ± 0.4 | 2.32 ± 0.3 |
|  | HHT | 49.50 ± 2.4 ↑ | 29.42 ± 1.8 ↑ | 13.59 ± 1.6 ↑ | 9.00 ± 1.3 | 4.37 ± 0.4 | 2.21 ± 0.3 ↓ |
| Fz | FFT | 43.89 ± 3.7 | 25.98 ± 1.6 | 12.53 ± 1.6 | 8.38 ± 0.9 | 4.17 ± 0.3 | 2.28 ± 0.3 |
|  | HHT | 50.00 ± 2.8 ↑ | 29.47 ± 1.9 ↑ | 13.77 ± 1.8 ↑ | 8.34 ± 0.9 | 4.19 ± 0.3 | 2.21 ± 0.3 ↓ |
| Cz | FFT | 43.22 ± 3.2 | 25.82 ± 1.8 | 12.33 ± 1.4 | 8.62 ± 0.9 | 4.27 ± 0.4 | 2.29 ± 0.3 |
|  | HHT | 49.40 ± 2.3 ↑ | 29.41 ± 1.9 ↑ | 13.50 ± 1.7 ↑ | 8.61 ± 0.9 | 4.29 ± 0.4 | 2.19 ± 0.3 ↓ |
| Pz | FFT | 43.06 ± 3.3 | 25.76 ± 1.7 | 12.21 ± 1.2 | 8.90 ± 1.0 | 4.36 ± 0.4 | 2.29 ± 0.3 |
|  | HHT | 49.35 ± 2.4 ↑ | 29.37 ± 2.0 ↑ | 13.36 ± 1.5 ↑ | 8.86 ± 1.1 | 4.37 ± 0.5 | 2.20 ± 0.3 ↓ |

Results are expressed as mean ± standard deviation; ↑\_ Significant higher, or ↓\_ lower values (p <0.05) for the ANOVA "F" statistics of values calculated using the FFT or the HHT methods.

**Table 5.** Power spectral density expressed in normalized units (%) calculated from the intrinsic mode functions extracted from the EEG of thirty-four healthy volunteers using two spectral methods.

| Leads | Method | IMF1 | IMF2 | IMF3 | IMF4 | IMF5 | IMF6 |
| --- | --- | --- | --- | --- | --- | --- | --- |
| Fp1 | FFT | 6.72 ± 6.3 | 13.41 ± 8.4 | 42.75 ± 15.4 | 22.40 ± 8.5 | 10.77 ± 5.4 | 3.94 ± 1.8 |
|  | HHT | 7.02 ± 6.3 | 13.57 ± 8.2 | 42.43 ± 15.2 | 22.19 ± 8.3 | 10.69 ± 5.4 ↓ | 4.11 ± 1.8 |
| Fp2 | FFT | 6.33 ± 4.8 | 13.22 ± 8.3 | 42.72 ± 15.3 | 22.44 ± 8.7 | 11.14 ± 5.7 | 4.15 ± 2.1 |
|  | HHT | 6.61 ± 4.9 | 13.39 ± 8.2 | 42.50 ± 15.2 | 22.19 ± 8.4 | 10.99 ± 5.5 | 4.32 ± 2.1 |
| F3 | FFT | 5.63 ± 3.8 | 13.60 ± 8.8 | 45.13 ± 15.7 | 22.82 ± 8.7 | 9.67 ± 4.7 | 3.16 ± 1.4 |
|  | HHT | 5.94 ± 4.0 | 13.79 ± 8.7 | 44.89 ± 15.4 | 22.54 ± 8.5 | 9.55 ± 4.7 | 3.29 ± 1.4 |
| F4 | FFT | 5.17 ± 3.3 | 13.37 ± 8.5 | 45.29 ± 15.8 | 22.67 ± 8.6 | 10.13 ± 5.3 | 3.38 ± 1.8 |
|  | HHT | 5.46 ± 3.5 | 13.59 ± 8.4 | 45.09 ± 15.6 | 22.34 ± 8.3 | 10.01 ± 5.2 | 3.50 ± 1.8 |
| C3 | FFT | 4.84 ± 4.4 | 12.56 ± 8.3 | 48.69 ± 14.4 | 22.85 ± 9.1 | 8.52 ± 4.8 | 2.55 ± 1.4 |
|  | HHT | 5.10 ± 4.5 | 12.81 ± 8.2 | 48.43 ± 14.3 | 22.61 ± 8.9 | 8.40 ± 4.7 | 2.65 ± 1.4 ↑ |
| C4 | FFT | 4.39 ± 3.6 | 12.10 ± 7.9 | 49.49 ± 13.6 | 22.85 ± 8.7 | 8.61 ± 4.9 | 2.56 ± 1.3 |
|  | HHT | 4.63 ± 3.8 | 12.32 ± 7.8 | 49.20 ± 13.6 | 22.66 ± 8.5 | 8.56 ± 4.9 | 2.63 ± 1.3 |
| P3 | FFT | 4.29 ± 6.8 | 9.90 ± 7.7 | 50.89 ± 14.7 | 24.96 ± 11.3 | 7.50 ± 4.8 | 2.45 ± 1.5 |
|  | HHT | 4.49 ± 6.9 | 10.14 ± 7.6 | 50.58 ± 14.7 | 24.76 ± 11.1 | 7.47 ± 4.8 | 2.56 ± 1.5 ↑ |
| P4 | FFT | 3.69 ± 3.9 | 10.30 ± 8.0 | 51.17 ± 13.6 | 24.46 ± 10.2 | 8.00 ± 4.6 | 2.38 ± 1.2 |
|  | HHT | 3.87 ± 4.0 | 10.49 ± 7.9 | 50.91 ± 13.7 | 24.30 ± 10.1 | 7.96 ± 4.6 | 2.48 ± 1.2 ↑ |
| O1 | FFT | 3.39 ± 3.3 | 8.67 ± 7.2 | 53.81 ± 14.9 | 24.82 ± 11.7 | 7.05 ± 4.1 | 2.27 ± 1.1 |
|  | HHT | 3.55 ± 3.4 ↑ | 8.82 ± 7.1 ↑ | 53.41 ± 14.8 | 24.80 ± 11.5 | 7.04 ± 4.1 | 2.38 ± 1.1 |
| O2 | FFT | 2.95 ± 2.9 | 7.90 ± 6.8 | 55.86 ± 15.4 | 24.8 ± 12.1 | 6.51 ± 3.8 | 1.99 ± 0.9 |
|  | HHT | 3.12 ± 3.1 ↑ | 8.04 ± 6.8 ↑ | 55.49 ± 15.3 ↓ | 24.82 ± 12.1 | 6.47 ± 3.8 ↓ | 2.07 ± 0.9 ↑ |
| F7 | FFT | 6.03 ± 4.8 | 12.89 ± 7.4 | 43.50 ± 14.8 | 23.48 ± 8.7 | 10.49 ± 5.1 | 3.62 ± 1.6 |
|  | HHT | 6.34 ± 5.0 | 13.10 ± 7.3 ↑ | 43.13 ± 14.5 | 23.22 ± 8.5 | 10.43 ± 5.1 | 3.79 ± 1.6 |
| F8 | FFT | 6.40 ± 5.1 | 12.71 ± 7.2 | 43.77 ± 14.5 | 22.53 ± 8.5 | 10.90 ± 5.4 | 3.69 ± 1.7 |
|  | HHT | 6.67 ± 5.2 | 12.91 ± 7.1 | 43.43 ± 14.4 | 22.31 ± 8.3 | 10.84 ± 5.2 ↑ | 3.85 ± 1.8 |
| T3 | FFT | 7.26 ± 5.7 | 13.57 ± 7.8 | 44.30 ± 13.9 | 23.40 ± 9.0 | 8.69 ± 4.6 | 2.76 ± 1.3 |
|  | HHT | 7.62 ± 5.9 | 13.81 ± 7.7 | 43.96 ± 13.7 | 23.09 ± 8.7 | 8.64 ± 4.6 | 2.87 ± 1.4 |
| T4 | FFT | 7.26 ± 6.0 | 13.83 ± 8.0 | 45.27 ± 13.2 | 22.21 ± 8.4 | 8.75 ± 5.0 | 2.68 ± 1.3 |
|  | HHT | 7.55 ± 6.2 | 14.02 ± 7.9 | 44.93 ± 13.3 | 22.02 ± 8.4 | 8.70 ± 4.9 | 2.77 ± 1.3 |
| T5 | FFT | 4.97 ± 3.9 | 10.69 ± 6.8 | 47.51 ± 13.3 | 26.31 ± 11.2 | 7.95 ± 4.6 | 2.57 ± 1.1 |
|  | HHT | 5.21 ± 4.1 | 10.91 ± 6.7 | 47.09 ± 13.3 | 26.19 ± 11.2 ↓ | 7.92 ± 4.5 | 2.69 ± 1.2 ↑ |
| T6 | FFT | 4.83 ± 4.9 | 10.65 ± 7.3 | 48.75 ± 14.6 | 25.39 ± 11.2 | 7.86 ± 4.8 | 2.52 ± 1.2 |
|  | HHT | 5.06 ± 5.0 | 10.84 ± 7.3 | 48.40 ± 14.6 | 25.25 ± 11.1 | 7.84 ± 4.8 | 2.61 ± 1.2 |
| Fz | FFT | 4.15 ± 2.7 | 12.18 ± 8.5 | 45.51 ± 16.6 | 23.97 ± 9.5 | 10.84 ± 5.8 | 3.36 ± 1.9 |
|  | HHT | 4.40 ± 2.8 | 12.35 ± 8.3 | 45.29 ± 16.3 | 23.75 ± 9.2 | 10.72 ± 5.7 | 3.50 ± 1.9 |
| Cz | FFT | 3.38 ± 2.7 | 12.38 ± 8.9 | 48.28 ± 15.1 | 23.48 ± 9.6 | 9.59 ± 5.4 | 2.88 ± 1.5 |
|  | HHT | 3.62 ± 2.9 | 12.59 ± 8.8 | 48.04 ± 15.0 | 23.29 ± 9.4 | 9.50 ± 5.4 | 2.97 ± 1.5 |
| Pz | FFT | 2.88 ± 2.8 | 10.41 ± 8.9 | 51.81 ± 15.1 | 24.21 ± 10.7 | 8.12 ± 5.1 | 2.57 ± 1.4 |
|  | HHT | 3.07 ± 2.9 ↑ | 10.62 ± 8.8 | 51.51 ± 14.9 | 24.05 ± 10.5 | 8.07 ± 5.0 | 2.67 ± 1.4 |

Results are expressed as mean ± standard deviation; ↑\_ Significant higher, or ↓\_ lower values (p <0.05) for the ANOVA “F” statistics of values calculated using the FFT or the HHT methods; original values required a *logit* transformation to achieve a normal distribution before the statistical testing.

The range for the mean weighted frequency values (mean ± 1.96*DS) for each EEG lead and IMF calculated with the two methods are shown in Table 6. Only was detected in some degree the overlapping of spectral frequencies between the IMF3 and IMF4 using this metric.

**Table 6.** Lower and upper limits for the range mean ± 1.96 DS of the mean weighted frequency values calculated for the intrinsic mode functions extracted from the EEG of thirty-four healthy volunteers using two spectral methods.

| Leads | Method | IMF1 | IMF2 | IMF3 | IMF4 | IMF5 | IMF6 |
| --- | --- | --- | --- | --- | --- | --- | --- |
| Fp1 | FFT | 36.9 – 51.8 | 22.6 – 29.5 | 9.6 – 15.5* | 6.8 – 10.2* | 3.5 – 5.0 | 1.8 – 2.8 |
|  | HHT | 44.2 – 55.5 | 25.5 – 32.9 | 10.3 – 17.0 | 6.7 – 10.3 | 3.4 – 5.1 | 1.6 – 2.8 |
| Fp2 | FFT | 36.5 – 52.0 | 22.6 – 29.7 | 9.6 – 15.4* | 6.6 – 10.2* | 3.5 – 4.9 | 1.8 – 2.7 |
|  | HHT | 44.0 – 55.6 | 25.8 – 32.8 | 10.3 – 16.8* | 6.4 – 10.4* | 3.5 – 5.0 | 1.6 – 2.7 |
| F3 | FFT | 36.7 – 51.2 | 22.2 – 29.7 | 9.5 – 15.5* | 6.8 – 10.2* | 3.7 – 4.9 | 1.8 – 2.7 |
|  | HHT | 44.5 – 54.8 | 25.3 – 33.2 | 10.4 – 16.9 | 6.4 – 10.4 | 3.6 – 5.0 | 1.6 – 2.8 |
| F4 | FFT | 37.1 – 50.4 | 22.4 – 29.5 | 9.6 – 15.4* | 6.8 – 10.1* | 3.5 – 4.9 | 1.8 – 2.7 |
|  | HHT | 44.9 – 54.1 | 25.7 – 32.9 | 10.3 – 17.1 | 6.6 – 10.1 | 3.5 – 5.0 | 1.7 – 2.7 |
| C3 | FFT | 36.2 – 50.8 | 22.2 – 29.4 | 9.6 – 15.0* | 7.0 – 10.3* | 3.5 – 5.1 | 1.7 – 2.8 |
|  | HHT | 44.5 – 54.5 | 25.6 – 33.2 | 10.3 – 16.6* | 6.8 – 10.6* | 3.5 – 5.2 | 1.6 – 2.8 |
| C4 | FFT | 35.9 – 51.5 | 21.9 – 29.7 | 9.8 – 14.7* | 6.9 – 10.5* | 3.6 – 5.0 | 1.8 – 2.7 |
|  | HHT | 44.3 – 55.3 | 25.3 – 33.4 | 10.5 – 16.4* | 6.7 – 10.7* | 3.6 – 5.0 | 1.7 – 2.7 |
| P3 | FFT | 35.7 – 51.2 | 22.3 – 29.2 | 9.8 – 14.9* | 7.2 – 10.8* | 3.6 – 5.1 | 1.8 – 2.7 |
|  | HHT | 44.4 – 54.8 | 25.7 – 33.0 | 10.3 – 16.7* | 6.8 – 11.2* | 3.6 – 5.1 | 1.7 – 2.7 |
| P4 | FFT | 36.0 – 51.0 | 22.2 – 29.1 | 9.9 – 14.7* | 7.0 – 10.9* | 3.5 – 5.1 | 1.7 – 2.9 |
|  | HHT | 44.6 – 54.6 | 25.5 – 33.2 | 10.6 – 16.5* | 6.7 – 11.3* | 3.6 – 5.0 | 1.6 – 2.8 |
| O1 | FFT | 36.6 – 50.7 | 22.4 – 28.9 | 9.7 – 14.5* | 7.0 – 11.2* | 3.5 – 5.1 | 1.9 – 2.8 |
|  | HHT | 45.1 – 54.0 | 25.9 – 32.7 | 10.2 – 16.4* | 6.7 – 11.6* | 3.6 – 5.1 | 1.7 – 2.7 |
| O2 | FFT | 37.2 – 49.7 | 22.6 – 29.0 | 9.5 – 14.5* | 7.1 – 11.5* | 3.6 – 5.2 | 1.8 – 2.8 |
|  | HHT | 45.9 – 52.7 | 26.1 – 32.8 | 10.0 – 16.3* | 6.7 – 12.0* | 3.5 – 5.1 | 1.6 – 2.9 |
| F7 | FFT | 37.5 – 50.2 | 22.4 – 29.4 | 9.6 – 15.4* | 6.7 – 10.4* | 3.6 – 4.9 | 1.9 – 2.7 |
|  | HHT | 36.8 – 51.3 | 22.3 – 29.7 | 9.7 – 15.2* | 6.7 – 10.4* | 3.5 – 5.0 | 1.9 – 2.8 |
| F8 | FFT | 37.4 – 50.3 | 22.3 – 29.8 | 9.9 – 15.2* | 6.8 – 10.5* | 3.6 – 5.0 | 1.9 – 2.7 |
|  | HHT | 37.0 – 50.8 | 22.0 – 30.0 | 9.9 – 15.0* | 6.7 – 10.7* | 3.5 – 5.1 | 1.8 – 2.9 |
| T3 | FFT | 37.4 – 50.3 | 22.3 – 29.8 | 9.9 – 15.2* | 6.8 – 10.5* | 3.6 – 5.0 | 1.9 – 2.7 |
|  | HHT | 44.9 – 53.9 | 25.8 – 32.7 | 10.5 – 16.8* | 6.5 – 10.8* | 3.6 – 5.0 | 1.7 – 2.7 |
| T4 | FFT | 37.0 – 50.8 | 22.0 – 30.0 | 9.9 – 15.0* | 6.7 – 10.7* | 3.5 – 5.1 | 1.8 – 2.9 |
|  | HHT | 45.1 – 53.9 | 25.8 – 32.7 | 10.8 – 16.3 | 6.5 – 10.8 | 3.6 – 5.0 | 1.6 – 2.8 |
| T5 | FFT | 36.4 – 51.5 | 22.4 – 29.3 | 9.9 – 14.9* | 7.0 – 10.9* | 3.8 – 4.9 | 1.8 – 2.8 |
|  | HHT | 44.5 – 54.9 | 26.0 – 32.3 | 10.5 – 16.6* | 6.6 – 11.4* | 3.7 – 4.9 | 1.7 – 2.7 |
| T6 | FFT | 36.6 – 50.8 | 22.5 – 29.3 | 9.8 – 15.0* | 6.8 – 11.1* | 3.6 – 5.2 | 1.8 – 2.8 |
|  | HHT | 44.8 – 54.2 | 26.0 – 32.9 | 10.4 – 16.8* | 6.4 – 11.6* | 3.7 – 5.1 | 1.6 – 2.8 |
| Fz | FFT | 36.6 – 51.2 | 22.8 – 29.2 | 9.4 – 15.6* | 6.7 – 10.1* | 3.5 – 4.9 | 1.8 – 2.8 |
|  | HHT | 44.5 – 55.5 | 25.7 – 33.3 | 10.3 – 17.3 | 6.5 – 10.2* | 3.5 – 4.9 | 1.6 – 2.8 |
| Cz | FFT | 37.0 – 49.4 | 22.3 – 29.3 | 9.5 – 15.2* | 6.9 – 10.4* | 3.5 – 5.0 | 1.8 – 2.8 |
|  | HHT | 44.9 – 53.8 | 25.6 – 33.2 | 10.2 – 16.8* | 6.8 – 10.4* | 3.5 – 5.1 | 1.7 – 2.7 |
| Pz | FFT | 36.6 – 49.5 | 22.4 – 29.1 | 9.8 – 14.6* | 6.9 – 10.9* | 3.5 – 5.2 | 1.8 – 2.8 |
|  | HHT | 44.7 – 54.0 | 25.5 – 33.3 | 10.3 – 16.4* | 6.7 – 11.0* | 3.5 – 5.3 | 1.6 – 2.8 |
| AVE | LFL | 40.29 ± 4.0 | 23.84 ± 1.7 | 10.01 ± 0.4 | 6.74 ± 0.2 | 3.56 ± 0.1 | 1.72 ± 0.1 |
| | UFL | 52.43 ± 2.0 | 31.03 ± 1.8 | 15.79 ± 0.9 | 10.71 ± 0.5 | 5.04 ± 0.1 | 2.77 ± 0.1 |

Results are expressed as lower limit – upper limit values for each EEG lead; AVE: Grand average value of the lower limits (LFL) and upper limits (UFL) including all EEG leads expressed as mean ± standard deviation; \*: Denotes the presence of some degree of overlapping with a near IMF

## 4. Discussion

The results obtained using the two methods in this study showed that the first 6 oscillatory component modes (IMFs) extracted from the EEG of the 19 explored scalp positions, contained frequencies that corresponded approximately to the range from 1.5 to 56 Hz, considering the observed mean weighted frequency ranges (Table 4). This range corresponds with the expected frequencies of the EEG from the classic delta to the gamma bands. The estimations of the power spectral density expressed in normalized units (%) of the oscillatory components (IMFs) didn’t show notable differences between methods (Table 3), and the detected significant differences of the mean weighted frequencies can be ascribed to the fact that the HHT method has a higher instantaneous frequency resolution than the FFT method as noted by (Durstewitz, Deco 2008; Munoz-Gutierrez et al. 2018), and the fact that Fourier transform have constant frequencies and weights, while the Hilbert transform allows the frequency as well as the amplitudes to vary over time (Al-Subari et al. 2015).

Some authors have recently suggested that the first IMFs decomposed from the EEG using the multivariate empirical mode decomposition could be associated to classical EEG bands (Noshadi et al. 2014; Schiecke et al. 2019; Soler et al. 2020; Tsai et al. 2016). The IMF1 would correspond to the gamma band, the IMF2 to the beta, the IMF3 to the alpha, the IMF4 to the theta, and the IMF5 and IMF6 would be sub-bands of the delta band (Tsai et al. 2016). Our results don’t agree completely with these. If we pay attention to the grand average ranges of the mean weighted frequencies shown in Table 6 we can conclude that the IMF1 showed for our group values from 40.3 - 52.4 Hz that correspond to the gamma band, for the IMF2 values of 23.84 – 31.03 corresponding to the beta-2 band, for the IMF5 values from 3.56 – 5.04 corresponding to the theta band, for the IMF6 from 1.72 – 2.77 corresponding to the delta band, but for the IMF3 values were observed from 10.01 – 15.79, including frequencies of the alpha and beta-1 band, while for the IMF4 the observed values were from 6.74 – 10.71 Hz that belong to the theta and alpha bands, and both frequency ranges (IMF3 and IMF4) were slightly overlapped. This behavior was observed for all the EEG leads (Table 6), and was also detected for the grand averages of the distribution of the mean weighted frequency (Figure 4 A), and the grand averages of the spectra of the IMFs in the group of subjects for the C3 EEG lead (Figure 4 B,C). However, the fact that the PSD values in normalized values (%) for the IMF3 were almost two times than those observed for the IMF4 (Table 5) allows us to speculate that the IMF3 component reflects better the alpha spectral frequencies, and could be considered as mainly related to this classic range, while the IMF4 could be more associated to the theta band.

These findings are the evidence of one of the limitations of the empirical mode decomposition: the so called mode-mixing, consisting in the appearance of disparate scales in an IMF, or when a signal with a similar scale appears in different IMF components. This fact has been reported and multiple procedures have been proposed to mitigate its effect (Munoz-Gutierrez et al. 2018; Soler et al. 2020; Tsai et al. 2016; Zheng, Xu 2019).

In the present study were applied different strategies to mitigate the mode-mixing. The EMD method that was applied to decompose the EEG was the most novel of the multivariate versions, the APIT-MEMD (Hemakom et al. 2016) that has been developed as an improvement to the MEMD (Rehman, Mandic 2010; Ur Rehman et al. 2010), with the express purpose to cope with power imbalances and inter-channel correlations of multichannel signals as the EEG signal, and also was used the method proposed by (Xie, Wang 2006) for the calculation of the mean spectral frequencies that mitigate the effect of some frequencies that have associated low values of power spectral density. However, as has been acknowledged by other authors the mode-mixing problem continues to be a big issue for the analysis of multichannel signals like the EEG (Alegre-Cortes et al. 2016; Munoz-Gutierrez et al. 2018; Soler et al. 2020; Zheng, Xu 2019).

However, it must be acknowledged also that different authors have questioned the limits generally established for the EEG bands considering their findings that several oscillatory patterns can be present simultaneously and can modulate one to another (Buzsaki 2006; Buzsaki, Watson 2012; Penttonen, Buzsaki 2003). Our results could be an evidence of this fact and can coexist with the effect of the mode-mixing, but it results impossible to separate them.

The two methods used in this study for the calculation of the investigated spectral indices proved to be equally informative. Although the HHT method requires an extra step in the processing, the application of the Hilbert transform to the IMFs, we recommend this approach because it produces a better frequency resolution, as a result of the properties of the Hilbert-Huang transform method.

Using both methods can be calculated indices for broad band or narrow band spectral analysis of the multicomponent oscillatory modes of the EEG that can become in the near future an alternative to the classic spectral analysis of the original EEG signals.

## 5. Conclusions

The decomposition of the EEG signals recorded simultaneously in healthy humans from several locations of the scalp using a multivariate empirical mode decomposition algorithm (APIT-MEMD in this study) allows to obtain different multicomponent oscillatory modes whose frequency ranges are clearly related with those ascribed to spectral bands studied in classical electroencephalography. The use of the FFT approach to obtain the spectra of these modes to study different indices calculated in the frequency domain (mean frequencies and power spectral density) results equally informative to the use of the Hilbert-Huang transform. However, the HHT method may be recommended considering its better frequency resolution. The overlapping of frequencies observed for some of the oscillatory modes, in our case for the third and fourth modes, related to the so called mode-mixing problem of the EMD algorithms, and reported by many other authors, was also observed in this study. This fact doesn’t denies that this overlapping of frequencies observed in our study between oscillatory modes can be also associated with the reported findings that several oscillatory patterns can be present simultaneously and can modulate one to another in the EEG. This novel approach for the study of EEG rhythms may become in the near future a clear alternative to the FFT method, considering the presence in the EEG signals of non-Gaussian, nonlinear, and non-stationary processes that may produce misleading results when using the FFT approach that strictly speaking is valid only for linear and stationary processes.

## Competing Interests

The authors declare that they have no competing interests in relation to this article.

## Notes

### Competing Interest Statement

The authors have declared no competing interest.

## References

Abdulhay E, Alafeef M, Abdelhay A, Al-Bashir A. Classification of Normal, Ictal and Inter-ictal EEG via Direct Quadrature and Random Forest Tree. Journal of medical and biological engineering. 2017;37(6):843–57. doi:10.1007/s40846-017-0239-z.

Al-Subari K, Al-Baddai S, Tome AM, Volberg G, Hammwohner R, Lang EW. Ensemble Empirical Mode Decomposition Analysis of EEG Data Collected during a Contour Integration Task. PloS one. 2015;10(4):e0119489. doi:10.1371/journal.pone.0119489.

Al-Subari K, Al-Baddai S, Tome AM, Volberg G, Ludwig B, Lang EW. Combined EMD-sLORETA Analysis of EEG Data Collected during a Contour Integration Task. PloS one. 2016;11(12):e0167957. doi:10.1371/journal.pone.0167957.

Alegre-Cortes J, Soto-Sanchez C, Piza AG, Albarracin AL, Farfan FD, Felice CJ et al. Time-frequency analysis of neuronal populations with instantaneous resolution based on noise-assisted multivariate empirical mode decomposition. Journal of neuroscience methods. 2016;267:35–44. doi:10.1016/j.jneumeth.2016.03.018.

Amo C, de Santiago L, Barea R, Lopez-Dorado A, Boquete L. Analysis of Gamma-Band Activity from Human EEG Using Empirical Mode Decomposition. Sensors. 2017;17(5). doi:10.3390/s17050989.

Berger H. Über das Elektrenkephalogramm des Menschen. Arch Psychiat Nervenkr. 1929;87:527–70.

Buzsaki G. Rhythms of the brain. Oxford University Press, Inc.; 2006.

Buzsaki G, Watson BO. Brain rhythms and neural syntax: implications for efficient coding of cognitive content and neuropsychiatric disease. Dialogues in clinical neuroscience. 2012;14(4):345–67.

Carella T, De Silvestri M, Finedore M, Haniff I, Esmailbeigi H. Emotion Recognition for Brain Machine Interface: Non-linear Spectral Analysis of EEG Signals Using Empirical Mode Decomposition. Conference proceedings: Annual International Conference of the IEEE Engineering in Medicine and Biology Society IEEE Engineering in Medicine and Biology Society Annual Conference. 2018;2018:223–6. doi:10.1109/EMBC.2018.8512228.

Cooley JW, Tukey JW. An algorithm for the machine calculation of complex Fourier series. Mathematics of Computation. 1965;19(90):297–301. doi:10.1090/S0025-5718-1965-0178586-1.

Chatterjee S. Detection of focal electroencephalogram signals using higher-order moments in EMD-TKEO domain. Healthc Technol Lett. 2019;6(3):64–9. doi:10.1049/htl.2018.5036.

Chen SJ, Peng CJ, Chen YC, Hwang YR, Lai YS, Fan SZ et al. Comparison of FFT and marginal spectra of EEG using empirical mode decomposition to monitor anesthesia. Computer methods and programs in biomedicine. 2016;137:77–85. doi:10.1016/j.cmpb.2016.08.024.

Chen YF, Atal K, Xie SQ, Liu Q. A new multivariate empirical mode decomposition method for improving the performance of SSVEP-based brain-computer interface. Journal of neural engineering. 2017;14(4):046028. doi:10.1088/1741-2552/aa6a23.

Chuang KY, Chen YH, Balachandran P, Liang WK, Juan CH. Revealing the Electrophysiological Correlates of Working Memory-Load Effects in Symmetry Span Task With HHT Method. Frontiers in psychology. 2019;10:855. doi:10.3389/fpsyg.2019.00855.

Dinares-Ferran J, Ortner R, Guger C, Sole-Casals J. A New Method to Generate Artificial Frames Using the Empirical Mode Decomposition for an EEG-Based Motor Imagery BCI. Frontiers in neuroscience. 2018;12:308. doi:10.3389/fnins.2018.00308.

Durstewitz D, Deco G. Computational significance of transient dynamics in cortical networks. Eur J Neurosci. 2008;27(1):217–27. doi:10.1111/j.1460-9568.2007.05976.x.

Estevez-Baez M, Machado C, Arrufat-Pie E, Santos A. Aplicación del método de Hilbert-Huang a señales biológicas en el campo de la neurología: descripción y aspectos metodológicos.. Technical Reports of the Institute of Neurology and Neurosurgery, Havana, Cuba. 2017a;Retrieved from https://www.researchgate.net/publication/320448562 (Date Accessed 15/12/2017):website: https://www.researchgate.net/.

Estevez-Baez M, Machado C, Arrufat-Pie E, Santos A. El método de Hilbert-Huang aplicado al estudio de algunas señales biológicas en el campo de la neurología: revisión bibliográfica. Technical Reports of the Institute of Neurology and Neurosurgery, Havana, Cuba. 2017b;Retrieved from https://www.researchgate.net/publication/320450700(Date accessed 15/12/2017):website: https://www.researchgate.net/.

Estevez-Baez M, Machado C, Arrufat-Pie E, Santos A. El método de Hilbert-Huang en el análisis del EEG: fundamentos y perspectivas.. Technical Reports of the Institute of Neurology and Neurosurgery, Havana, Cuba 2017c; Retrieved from https://www.researchgate.net/publication/320471970 (Date Accessed 15/12/2017):website: https://www.researchgate.net/.

Estevez-Baez M, Machado C, Garcia-Sanchez B, Rodriguez V, Alvarez-Santana R, Leisman G et al. Autonomic impairment of patients in coma with different Glasgow coma score assessed with heart rate variability. Brain injury. 2019;33(4):496–516. doi:10.1080/02699052.2018.1553312.

Estevez-Baez M, Machado C, Montes-Brown J, Jas-Garcia J, Leisman G, Schiavi A et al. Very High Frequency Oscillations of Heart Rate Variability in Healthy Humans and in Patients with Cardiovascular Autonomic Neuropathy. Advances in experimental medicine and biology. 2018;1070:49–70. doi:10.1007/5584_2018_154.

Hansen ST, Hemakom A, Gylling Safeldt M, Krohne LK, Madsen KH, Siebner HR et al. Unmixing Oscillatory Brain Activity by EEG Source Localization and Empirical Mode Decomposition. Computational intelligence and neuroscience. 2019;2019:5618303. doi:10.1155/2019/5618303.

Hassan AR, Bhuiyan MIH. Automated identification of sleep states from EEG signals by means of ensemble empirical mode decomposition and random under sampling boosting. Computer methods and programs in biomedicine. 2017;140:201–10. doi:10.1016/j.cmpb.2016.12.015.

Hemakom A, Goverdovsky V, Looney D, Mandic DP. Adaptive-projection intrinsically transformed multivariate empirical mode decomposition in cooperative brain-computer interface applications. Philosophical transactions Series A, Mathematical, physical, and engineering sciences. 2016;374(2065):20150199. doi:10.1098/rsta.2015.0199.

Hou F, Yu Z, Peng CK, Yang A, Wu C, Ma Y. Complexity of Wake Electroencephalography Correlates With Slow Wave Activity After Sleep Onset. Frontiers in neuroscience. 2018;12:809. doi:10.3389/fnins.2018.00809.

Huang NE, Shen Z, Long SR, Wu MC, Shih HH, Zheng Q et al. The empirical mode decomposition and the Hilbert spectrum for nonlinear and non-stationary time series analysis. Proc R Soc Lond. 1998;454:903–95. doi:10.1098/rspa.1998.0193.

Hughes SW, Lorincz ML, Parri HR, Crunelli V. Infraslow (<0.1 Hz) oscillations in thalamic relay nuclei basic mechanisms and significance to health and disease states. Progress in brain research. 2011;193:145–62. doi:10.1016/B978-0-444-53839-0.00010-7.

Javed E, Croce P, Zappasodi F, Gratta CD. Hilbert Spectral Analysis of EEG Data reveals Spectral Dynamics associated with Microstates. Journal of neuroscience methods. 2019:108317. doi:10.1016/j.jneumeth.2019.108317.

Malekzadeh-Arasteh O, Pu H, Lim J, Liu CY, Do AH, Nenadic Z et al. An Energy-Efficient CMOS Dual-Mode Array Architecture for High-Density ECoG-Based Brain-Machine Interfaces. IEEE transactions on biomedical circuits and systems. 2020;14(2):332–42. doi:10.1109/TBCAS.2019.2963302.

Munoz-Gutierrez PA, Giraldo E, Bueno-Lopez M, Molinas M. Localization of Active Brain Sources From EEG Signals Using Empirical Mode Decomposition: A Comparative Study. Frontiers in integrative neuroscience. 2018;12:55. doi:10.3389/fnint.2018.00055.

Noshadi S, Vahid Abootalebi V, Taghi Sadeghi M, Shahab Shahvazian M. Selection of an efficient feature space for EEG-based mental task discrimination. Biocybernetics and Biomedical Engineering 2014;34:159–68. doi:10.1016/j.bbe.2014.03.004.

Park C, Plank M, Snider J, Kim S, Huang HC, Gepshtein S et al. EEG gamma band oscillations differentiate the planning of spatially directed movements of the arm versus eye: multivariate empirical mode decomposition analysis. IEEE transactions on neural systems and rehabilitation engineering: a publication of the IEEE Engineering in Medicine and Biology Society. 2014;22(5):1083–96. doi:10.1109/TNSRE.2014.2332450.

Penttonen M, Buzsaki G. Natural logarithmic relationship between brain oscillators. Thalamus and Related Systems. 2003;2:145–52. doi:10.1016/S1472-9288(03)00007-4.

Rahman MM, Fattah SA. Mental Task Classification Scheme Utilizing Correlation Coefficient Extracted from Interchannel Intrinsic Mode Function. BioMed research international. 2017;2017:3720589. doi:10.1155/2017/3720589.

Rehman N, Mandic DP. Multivariate empirical mode decomposition. Proceedings of the Royal Society A. 2010;466(2117):1291–302. doi:10.1098/rspa.2009.0502.

Schiecke K, Schmidt C, Piper D, Putsche P, Feucht M, Witte H et al. Assignment of Empirical Mode Decomposition Components and Its Application to Biomedical Signals. Methods of information in medicine. 2015;54(5):461–73. doi:10.3414/ME14-02-0024.

Schiecke K, Schumann A, Benninger F, Feucht M, Baer K, Schlattmann P. Brain-Heart interactions considering complex physiological data: processing schemes for timevariant, frequency-dependent, topographical and statistical examination of directed interactions by Convergent Cross Mapping. Physiol Meas. 2019. doi:10.1088/1361-6579/ab5050.

Smyk MK, van Luijtelaar G. Circadian Rhythms and Epilepsy: A Suitable Case for Absence Epilepsy. Frontiers in neurology. 2020;11:245. doi:10.3389/fneur.2020.00245.

Soler A, Munoz-Gutierrez PA, Bueno-Lopez M, Giraldo E, Molinas M. Low-Density EEG for Neural Activity Reconstruction Using Multivariate Empirical Mode Decomposition. Frontiers in neuroscience. 2020;14:175. doi:10.3389/fnins.2020.00175.

Soltani Zangbar H, Ghadiri T, Seyedi Vafaee M, Ebrahimi Kalan A, Fallahi S, Ghorbani M et al. Theta Oscillations Through Hippocampal/Prefrontal Pathway: Importance in Cognitive Performances. Brain connectivity. 2020;10(4):157–69. doi:10.1089/brain.2019.0733.

Tsai FF, Fan SZ, Lin YS, Huang NE, Yeh JR. Investigating Power Density and the Degree of Nonlinearity in Intrinsic Components of Anesthesia EEG by the Hilbert-Huang Transform: An Example Using Ketamine and Alfentanil. PloS one. 2016;11(12):e0168108. doi:10.1371/journal.pone.0168108.

Ur Rehman N, Xia Y, Mandic DP. Application of multivariate empirical mode decomposition for seizure detection in EEG signals. Conference proceedings: Annual International Conference of the IEEE Engineering in Medicine and Biology Society IEEE Engineering in Medicine and Biology Society Annual Conference. 2010;2010:1650–3. doi:10.1109/IEMBS.2010.5626665.

van Putten MJ, Tjepkema-Cloostermans MC, Hofmeijer J. Infraslow EEG activity modulates cortical excitability in postanoxic encephalopathy. Journal of neurophysiology. 2015;113(9):3256–67. doi:10.1152/jn.00714.2014.

Vanhatalo S, Voipio J, Kaila K. Full-band EEG (FbEEG): an emerging standard in electroencephalography. Clin Neurophysiol. 2005;116(1):1–8. doi:S1388-2457(04)00374-8 [pii] 10.1016/j.clinph.2004.09.015.

Xie H, Wang Z. Mean frequency derived via Hilbert-Huang transform with application to fatigue EMG signal analysis. Computer methods and programs in biomedicine. 2006;82(2):114–20. doi:10.1016/j.cmpb.2006.02.009.

Zheng Y, Xu G. Quantifying mode mixing and leakage in multivariate empirical mode decomposition and application in motor imagery-based brain-computer interface system. Medical & biological engineering & computing. 2019;57(6):1297–311. doi:10.1007/s11517-019-01960-9.

Zhuang N, Zeng Y, Tong L, Zhang C, Zhang H, Yan B. Emotion Recognition from EEG Signals Using Multidimensional Information in EMD Domain. BioMed research international. 2017;2017:8317357. doi:10.1155/2017/8317357.

